# Determination of metal ion transport rate of human ZIP4 using stable zinc isotopes

**DOI:** 10.1101/2024.05.20.594990

**Authors:** Yuhan Jiang, Keith MacRenaris, Thomas V. O’Halloran, Jian Hu

## Abstract

The essential microelement zinc is absorbed in the small intestine mainly by the zinc transporter ZIP4, a representative member of the Zrt/Irt-like protein (ZIP) family. ZIP4 is reportedly upregulated in many cancers, making it a promising oncology drug target. To date, there have been no reports on the turnover number of ZIP4, which is a crucial missing piece of information needed to better understand the transport mechanism. In this work, we used a non-radioactive zinc isotope, ^70^Zn, and inductively coupled plasma mass spectrometry (ICP-MS) to study human ZIP4 (hZIP4) expressed in HEK293 cells. Our data showed that ^70^Zn can replace the radioactive ^65^Zn as a tracer in kinetic evaluation of hZIP4 activity. This approach, combined with the quantification of the cell surface expression of hZIP4 using biotinylation or surface-bound antibody, allowed us to estimate the apparent turnover number of hZIP4 to be in the range of 0.08-0.2 s^-1^. The turnover numbers of the truncated hZIP4 variants are significantly smaller than that of the full-length hZIP4, confirming a crucial role for the extracellular domain in zinc transport. Using ^64^Zn and ^70^Zn, we measured zinc efflux during the cell-based transport assay and found that it has little effect on the zinc import analysis under these conditions. Finally, we demonstrated that use of laser ablation (LA) ICP-TOF-MS on samples applied to a solid substrate significantly increased the throughput of the transport assay. We envision that the approach reported here can be applied to the studies of metal transporters beyond the ZIP family.

## Introduction

As an essential trace element, zinc ions play catalytic, structural, and regulatory roles in biological systems (1), including cell signaling (2-4) and cell cycle control during meiosis and mitosis (5-12). Zinc fluxes are also associated with numerous physio-pathological processes (3,5,9,13-19). In humans, dietary zinc is absorbed in the small intestine where its transport is primarily mediated by ZIP4, a zinc transporter belonging to the Zrt/Irt-like protein (ZIP) metal transporter family (20-23). The pivotal role of ZIP4 in zinc uptake is manifested by ZIP4 dysfunctional mutations that lead to acrodermatitis enteropathica (AE), a life-threatening recessive genetic disorder (24-28). Remarkably, aberrant expression of ZIP4 has been found in many types of cancer and knockdown of ZIP4 in cancer cells significantly reduces cell proliferation and metastasis (29-38). Limited tissue distribution under normal conditions and aberrant upregulation in cancer suggest that ZIP4 is a promising cancer biomarker and anti-tumor drug target.

The mechanism of zinc ion transport by ZIPs has not well understood. Recent structural and biochemical studies of a bacterial ZIP from *Bordetella bronchiseptica* have suggested that the ZIP transporters use an elevator mode as the common transport mechanism (39,40) (**Scheme S1A**). As elevator transporters can act either as carriers or channels to achieve substrate translocation across the membrane, it is important to determine the transport mechanism of the ZIPs of interest by characterizing their kinetic properties. Proteoliposome-based studies showed that the bacterial ZIPs may behave as carriers or ion channels (41-43), but the cell-based assays have suggested that the ZIPs act as carriers. Early studies on a ZIP homolog from *E*.*coli*, ZupT, showed that the transporter behaved as a carrier in an *E*.*coli* strain where several endogenous metal transporters have been knocked out (44,45). Due to the difficulties in protein production, liposome reconstitution, and the transporters’ sensitivity to lipid composition (46), eukaryotic ZIPs have only been studied in the cell-based transport assay (47-58). However, one concern with the cell-based assay is that metal efflux during the transport assay, which is due to the activities of metal exporters on the plasma membrane, may lead to an underestimation of the imported metal substrate, resulting in a distorted dose curve and incorrectly determined kinetic parameters. Therefore, it is necessary to evaluate the effect of metal efflux on the transport assay.

So far, the radioactive ^65^Zn is the most commonly used substrate in the cell-based transport assays for the ZIPs because of the high intracellular zinc level in mammalian cells (up to 250 μM). However, the inclusion of radioactive metals in the transport assay limits the application of this approach to the extent necessary to thoroughly characterize these important metal transporters. An alternative approach is to use a low-abundance stable isotope of zinc so that the high natural background issue can be largely relieved. Indeed, ^70^Zn, which takes only 0.6% of naturally present zinc, has been used in the cell-based transport assay (59), demonstrating the potential to study transport kinetics of metal transporters.

In this work, we used a modified ^70^Zn-based approach to study the transport kinetics of human ZIP4 (hZIP4) which is expressed in HEK293 cells. Quantification of ^70^Zn by ICP-MS and the cell surface expression level of the transporter allowed us to report, for the first time, the turnover number of hZIP4. Using this approach, we re-examined the role of the extracellular domain (ECD) for ZIP4 function and confirmed the crucial role of the ECD for optimal zinc transport. Using two stable zinc isotopes, ^64^Zn and ^70^Zn, we found that although zinc efflux is detectable it has little effect on the results of the transport assay under our experimental conditions. Taken together, these results lead us to conclude that hZIP4 most likely functions as a carrier, rather than an ion channel. We also demonstrated that the application of laser ablation (LA) ICP-MS can significantly increase the throughput of isotope measurement to study the cell samples loaded on a glass slide.

## Results and Discussion

### Preparation of the ^70^Zn-enriched culture media for the cell-based zinc transport assay

Due to the high concentration of zinc in mammalian cells (1), it is critical to distinguish between zinc imported from the extracellular space (exogenous zinc) and that already present in the cells prior to the assay (endogenous zinc). The use of the radioactive zinc isotope ^65^Zn is a well-established solution, but this long-lived and high-energy gamma emitter also raises safety concerns for the researchers performing the experiments. Therefore, the use of a non-radioactive metal substrate becomes an attractive option. In an early study, transport assays employed the stable isotope ^70^Zn (natural abundance=0.6%) (59). The ^70^Zn-enriched culture media was prepared in two steps: total zinc concentration in the culture media was first decreased by treatment with the zinc-specific S100A12 protein immobilized on agarose followed by supplementation with salts containing ^70^Zn (60). It was shown that the cellular ^70^Zn concentration was significantly increased after the cells were incubated with the ^70^Zn-enriched media and that the cells expressing ZIP transporters accumulated more ^70^Zn in a two-hour incubation than control cells, providing the proof-of-concept of using ^70^Zn in the cell-based transport assay.

Using an analogous approach, we treated the cell culture media containing fetal bovine serum (FBS) with the metal ion chelating resin Chelex-100 to lower the zinc concentration. Chelex-100 has relatively high selectivity toward zinc ions over the transition metals (manganese, iron, cobalt, nickel and copper) (61) and has been employed to prepare metal-deficient mammalian cell culture media (62). We tested whether a treatment of a commonly used mammalian culture media (DMEM+10% FBS) with Chelex-100 would be sufficient to remove most of the naturally present zinc so that the natural background of zinc would be low enough to allow for a sensitive analysis. As shown in **Table 1**, the results of inductively coupled plasma mass spectrometry (ICP-MS) analysis indicated that the Chelex-100 treatment removed approximately 90-95% of the zinc naturally present in the culture media. We then added the desired amounts of ^70^Zn (72% enrichment) to the Chelex-treated media to generate the ^70^Zn-enriched media for the cell-based transport assay. Consistent with the previous reports (60,62), the Chelex-treated media contain 43% less copper, 99% less calcium, 99% less magnesium when compared with the media without the treatment. These changes in metal content did not affect the activity of hZIP4 (**Figure S1**).

**Table 1.**
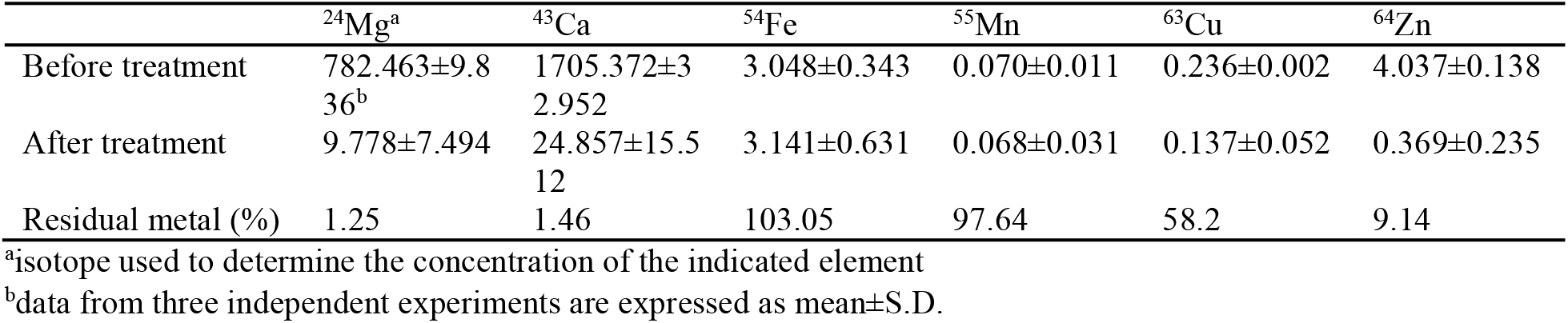
Changes of divalent metal concentrations (in μM) in the cell culture media before and after Chelex-100 treatment.

### Zinc transport assay of hZIP4 using the ^70^Zn-enriched culture media

To measure the transport activity of hZIP4, the ^70^Zn-enriched culture media was applied to HEK293 cells transiently expressing hZIP4. After incubation at 37 °C, the reaction was terminated with an ice-cold buffer containing 1 mM EDTA, followed by extensive wash using a phosphorous-free wash buffer. Concentrated nitric acid was then added to digest the cells and the resulting samples were subjected to ICP-MS analysis. To control for the variation in cell number between samples, we normalized the counts per second of ^70^Zn using the count ratio of ^70^Zn and ^31^P (^70^Zn/P) following the approach of Lane et al. (63). This calibration simplifies the procedure as there is no need to count cell numbers or measure protein concentrations of the samples. As shown in **Figure 1A**, incubation with the ^70^Zn-enriched culture media resulted in a higher level of the ^70^Zn/P ratio for the cells expressing hZIP4 than that for the cells transfected with an empty vector, indicating that the activity of hZIP4 can be tracked using the stable isotope ^70^Zn. Based on the time course experiments, an incubation time of 30 minutes was chosen for later experiments (**Figure 1B**).

**Figure 1.**
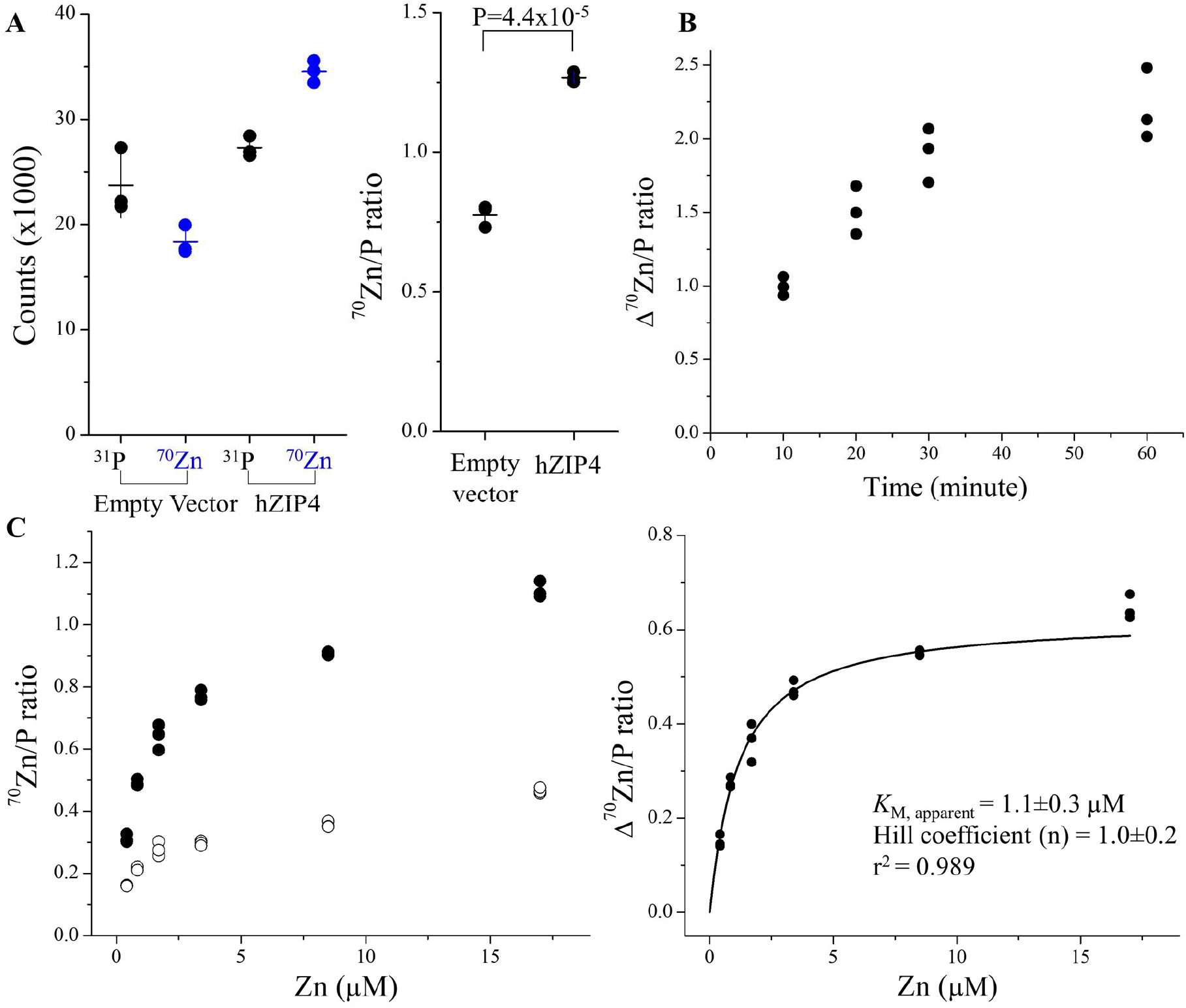
Cell-based kinetic study of hZIP4 using the stable isotope ^70^Zn as the substrate. (**A**) Counts (per second, detected in ICP-MS) of ^31^P and ^70^Zn in the cells with and without expressed hZIP4 (*left*) and the ^70^Zn/P count ratios (*right*). The error bars represent the standard deviation of the mean. The Student’s *t* test (two tailed) was performed to examine the statistically significant difference. (**B**) Time course of hZIP4-mediated ^70^Zn transport. Three replicates were performed at each time point. (**C**) Dose curve of hZIP4-mediated ^70^Zn transport. *Left*: The ^70^Zn/P count ratios of the cells with (solid circle) and without (open circle) expressing hZIP4 at the indicated concentrations of total zinc in the extracellular space. *Right*: Kinetic analysis of hZIP4. The activity of hZIP4 was calculated by subtracting the ^70^Zn/P count ratio of the cells transfected with the empty vector from that of the cells expressing hZIP4. The curve was fitted with the Hill model. Three replicates were performed for each condition. The extracellular zinc consists of ^70^Zn (with an enrichment of 72%) and other zinc isotopes. The shown data are from one of three independent experiments in which similar results were obtained.

Since the ^70^Zn/P ratios after background subtraction are linearly correlated with the cell surface expression levels of hZIP4 (**Figure S2**), the calculated ^70^Zn/P ratio represents the transport activity of hZIP4 expressed in HEK293T cells. Under the optimized conditions, a dose-dependent experiment was performed and the dose curve was fitted with the Hill model commonly used in steady-state kinetic study (**Figure 1C**). The apparent *K*_M_ is 1.1±0.3 μM with the Hill coefficient of ∼1, which agrees with our previously reported values when ^65^Zn was used in the transport assay (54,55).

### Determination of the turnover number of hZIP4 and re-examination of the role of the ECD

While the turnover number is a useful gauge for understanding transport mechanisms, there are no reports of such values to date for eukaryotic ZIP transporters. The turnover number of a transporter is commonly evaluated using experimentally determined values for *V*_max_ and the number of transporters expressed at the cell surface. *V*_max_ can be determined in the ICP-MS experiment that reports the total transport rate (number of zinc ions per second) of hZIP4 when zinc concentration is saturating (**Figure 1**) (54,56). To determine the cell surface expression level of a membrane protein, biotinylation followed by Western blot is a commonly used approach (64,65). We applied this approach to the cells expressing hZIP4-HA and estimated the number of hZIP4-HA expressed at the cell surface by comparison with the purified hZIP4 tagged with an N-FLAG and C-HA (FLAG-hZIP4-HA) in the Western blot experiment (**Figure 2A**). We also performed the ^70^Zn transport assay on the hZIP4-HA-expressing cells which were transfected at the same time as the cells used for the analysis of the hZIP4-HA surface expression (**Figure 2B**). Although the expression levels of hZIP4-HA may vary in different independent experiments, hZIP4-HA expression in the same batch of transfection are very similar (**Figure S3**). These results allowed us to estimate the apparent turnover number of hZIP4 to be 0.08±0.02 s^-1^ (mean±standard error, S.E., n=3). This value is generally consistent with the turnover numbers of representative carrier proteins (66) and close to a panel of slow carriers, including VMAT1 (0.2 s^-1^) (67), MATE1 (0.4 s^-1^) (68), NET (0.11 s^-1^) (69), Tyt1 (0.4 s^-1^) (70), and Glt_ph_ (0.14 s^-1^) (71). The slow transport rate and the saturable kinetics both suggest that hZIP4 is best characterized as a carrier. However, caution must be exercised because a low turnover number does not necessarily exclude the possibility of an ion channel, as the boundary between the two classes of membrane transport proteins can be blurred (72). Considering that ZIP4 undergoes endocytosis (73-75), it is plausible that the ZIP4 surface expression level determined by biotinylation may be overestimated. However, given that the zinc ion concentration in the PBS buffer used in the biotinylation reaction is unlikely to be at micromolar or higher and that the reaction was conducted only at room temperature for 10 minutes, we don’t think ZIP4 endocytosis, which is a zinc and energy dependent process, would significantly affect the estimation of ZIP4 expression at the cell surface.

**Figure 2.**
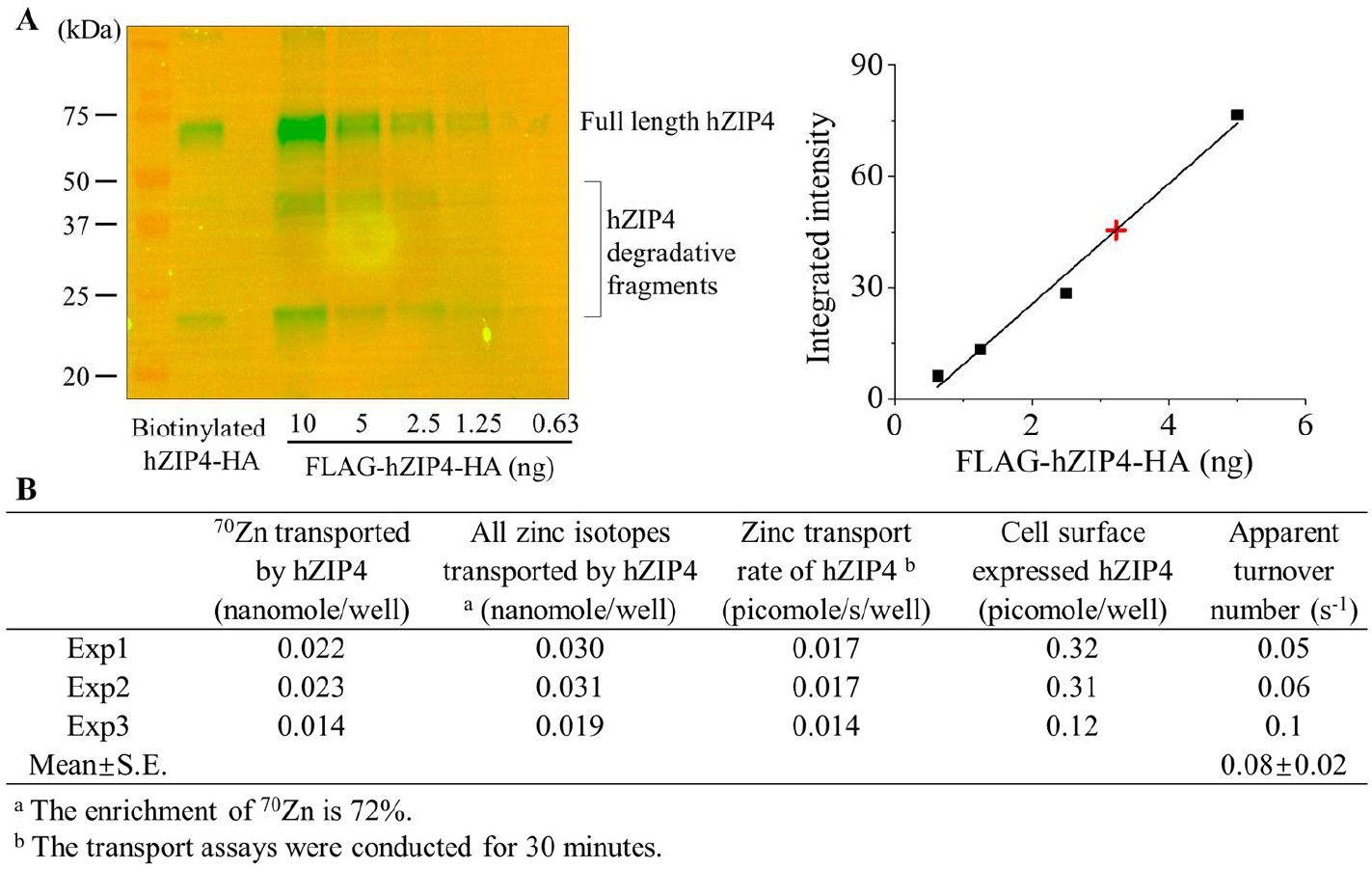
Determination of the turnover number of hZIP4. (**A**) Estimation of the surface expression level of hZIP4-HA by biotinylation and Western blot. The purified FLAG-hZIP4-HA was used as the standard. Degradative fragments were found in both biotinylated hZIP4-HA and FLAG-hZIP4-HA. In the standard curve, the black squares represent the standards and the red cross represents the biotinylated hZIP4-HA in the sample. Biotinylated sample was loaded into the gel with 50% dilution. The shown data are from one of three independent experiments. (**B**) Calculation of the turnover number of hZIP4 from three independent experiments.

We then extended this ^70^Zn-based assay to further evaluate the role of ZIP4-ECD in zinc transport function. ZIP4 has a large ECD among human ZIPs but the exact function of this domain has not been fully elucidated. In our early study where ^65^Zn was used in transport assays (54), we showed that deletion of the N-terminal-most helix-rich domain (HRD) resulted in a 50% loss of activity and deletion of the entire ECD, which consists of the HRD and the PAL-motif containing domain (PCD) (**Scheme S1** and **Figure S4**), resulted in approximately 75% loss of activity compared to the full-length hZIP4, indicating that the ECD is important for optimal zinc transport. However, a recent study reported that ECD deletion of hZIP4 with a C-terminal eGFP tag did not significantly affect the zinc transport activity when compared with the full-length hZIP4-eGFP (76). To clarify this point, we compared the apparent turnover numbers of hZIP4 variants with a C-terminal HA tag including the full-length transporter (hZIP4-HA), the ΔHRD variant (ΔHRD-hZIP4-HA), and the ΔECD variant (ΔECD-hZIP4-HA). The previous study has shown that the truncated variants did not affect the apparent *K*_M_ (54), so we focused only on the turnover number. The number of moles of ^70^Zn associated with the cells expressing hZIP4-HA or the variants were quantified by using ICP-MS as described above. In this study, we did not use biotinylation to estimate the cell surface expression levels of these constructs because they have different numbers of the lysine residues at the extracellular side and accessibility of the lysines differs according to a structure model predicted by AlphaFold (**Figure S4**). Instead, we determined the surface expression levels of hZIP4-HA and its variants by quantifying the anti-HA antibody bound at the surface of fixed but non-permeabilized cells. The use of a surface-bound antibody to evaluate the cell surface expression of a membrane protein is a well-established approach (77) and has been used in prior studies of ZIP4 (27,78). To quantify the cell surface-bound anti-HA antibodies, a standard curve was generated by applying the serially diluted anti-HA antibodies in a Western blot experiment **(Figure 3A)**. After calibrating the transport data with the cell surface expression levels of the HA-tagged transporter constructs (**Figure 3B**), the determined apparent turnover number of hZIP4-HA (0.2±0.03 s^-1^, mean±S.E., n=3, **Figure 3C**) was found to be generally consistent with that obtained from the biotinylation experiment (0.08±0.02 s^-1^, **Figure 2**). Consistent with our previous report (54), the cell surface expression levels of the two truncated variants are significantly higher than that of the full-length hZIP4-HA and the calculated apparent turnover numbers (0.05±0.01 s^-1^ for ΔHRD-hZIP4-HA, mean±S.E., n=3, and 0.03±0.01 s^-1^ for ΔECD-hZIP4-HA, mean±S.E., n=3) are significantly lower than that of the full-length hZIP4-HA. The difference in cell surface expression levels also agree with the different total expression levels for these constructs (**Figure S5**). Collectively, these results reinforce the notion that ZIP4-ECD is crucial for optimal zinc transport activity of hZIP4 and the HRD subdomain plays a more important role in supporting an efficient zinc transport than the PCD subdomain.

**Figure 3.**
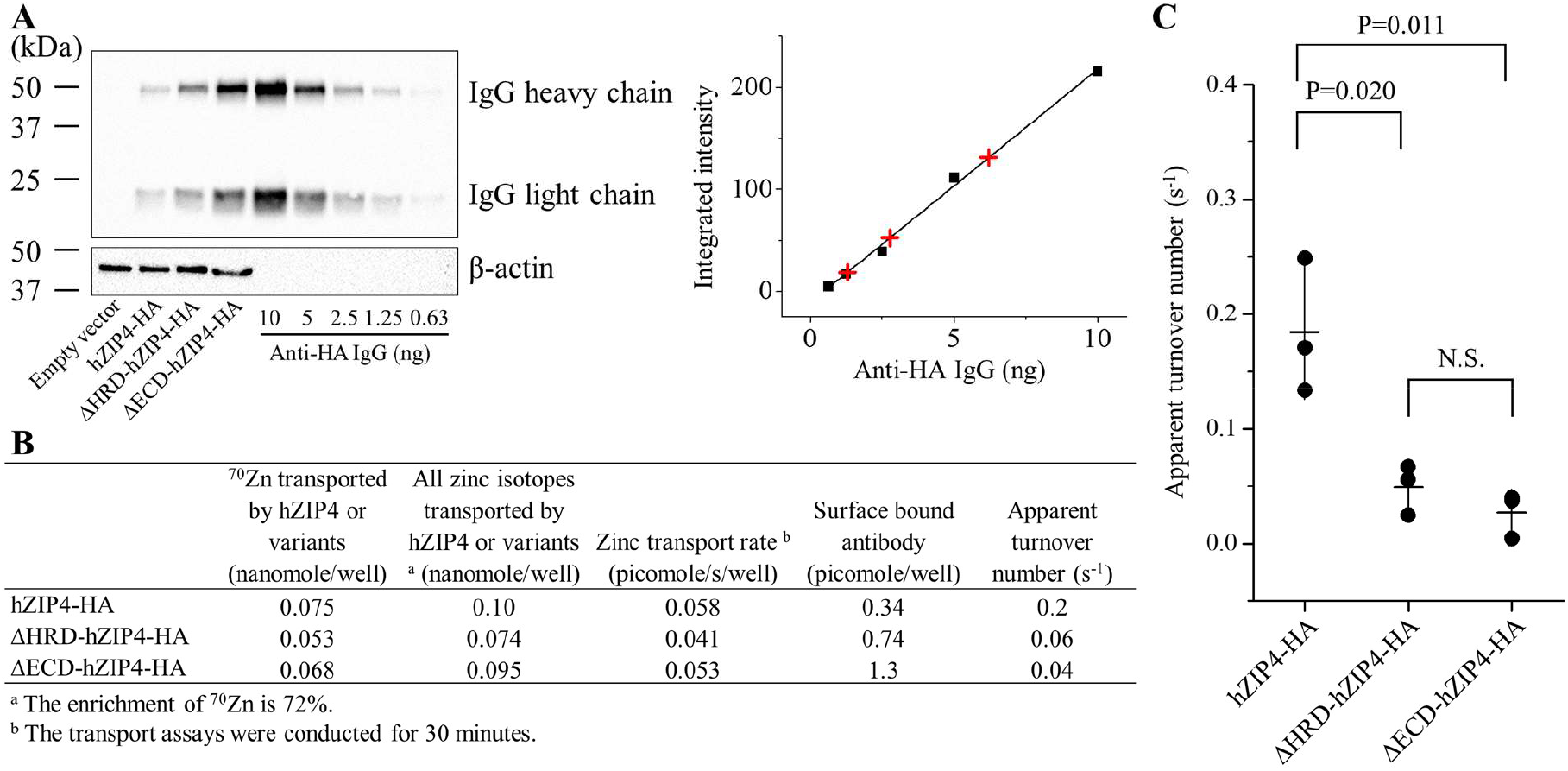
Comparison of the apparent turnover numbers of hZIP4-HA and the truncated variants (ΔHRD-hZIP4-HA and ΔECD-hZIP4-HA). (**A**) Estimation of the cell surface bound anti-HA antibodies. *Left*: Western blot of the cell surface bound anti-HA antibodies. The samples containing the indicated amount of anti-HA antibody were used as the standards. A HRP-conjugated horse-anti-mouse antibody was used to detect the mouse anti-HA antibodies. *Right*: Quantification of the cell surface bound anti-HA antibodies in the cell samples. The black squares represent the standards and the red plus signs represent the surface bound anti-HA antibody in the samples. (**B**) Calculation of the apparent turnover numbers using the data from one representative experiment. The shown data are from one of three independent experiments in which similar results were obtained. (**C**) Comparison of the apparent turnover numbers of hZIP4-HA, ΔHRD-hZIP4-HA, and ΔECD-hZIP4-HA. Each data point represents the mean of one of three independent experiments. Three replicates were performed in one experiment. The error bars represent the standard deviations of the mean. The Student’s *t* test (two tailed) was performed to examine statistically significant difference.

The exact function of ZIP4-ECD is still not fully understood. It has been shown that elimination of N-glycosylation of ZIP4-ECD does not affect the zinc transport activity (28). ZIP4-ECD forms a homodimer in the crystal structure primarily through the PCD subdomain (54). Although facilitating dimerization is a likely function of ZIP4-ECD, the fact that the HRD contributes most to promoting zinc transport (**Figure 3C**) suggests that the ECD must have additional function(s) beyond dimerization. It has been noted that all experimentally solved structures of BbZIP are in the inward-facing conformation, regardless of whether the transporter is in the substrate-bound or apo state (39,40,79), suggesting that the transporter in the outward-facing conformation is an energetically unfavorable state with a small population and that promoting the formation and/or stabilization of the outward-facing conformation may therefore represent a strategy to increase the transport activity. According to the structural model of ZIP4 and the proposed elevator mechanism (21,39,40), the HRD, which shows considerable flexibility in the crystal structure of ZIP4-ECD (54) and in the AlphaFold predicted structures (**Scheme S1B**), could transiently interact with the transport domain as the latter slides outward, thereby facilitating the formation and stabilization of the outward-facing conformation (**Scheme S1C**). Testing this hypothesis, especially when the structure of full-length ZIP4 is available, will help to clarify the function of the HRD that is present only in ZIP4 and its close homologue ZIP12.

### Estimation of zinc efflux during the transport assay

One concern with the cell-based transport assay is that while substrate is being transported into the cells by the importer of interest, some of the imported substrate may be expelled from the cells due to the activity of exporters at the plasma membrane. If this happens, it will lead to an underestimation of the influx rate and thus inaccuracy in the determination of the kinetic parameters. To evaluate possible effects of zinc efflux on the zinc transport assay for hZIP4, we designed the isotope exchange experiment. First, we determined how long it takes to replace the naturally present cellular zinc (endogenous zinc) with ^64^Zn (98% enrichment) added in the extracellular space. To do this, the cells growing in the regular culture media (DMEM plus 10% FBS, contains the naturally present zinc isotopes with natural abundance) were incubated with the ^64^Zn-enriched culture media and the levels of cellular ^70^Zn (as an indicator of the endogenous zinc in cells) and ^64^Zn were measured using ICP-MS at different time points. As shown in **Figure 4A**, after incubation for 48 hours without culture media replacement, the replacement of ^70^Zn by ^64^Zn reached equilibrium. The decrease in the ^70^Zn/^64^Zn count ratio fit well with the one-phase decay model, indicating that the imported ^64^Zn exchanged with a single intracellular zinc pool. Next, we cultured the cells in the ^64^Zn-enriched media for three passages (3 days per passage) to ensure a complete replacement and then transferred the cells to the same ^70^Zn-enriched media used in the zinc transport assay for a 30-min incubation. After termination by addition of EDTA followed by extensive washing, the amounts of ^64^Zn and ^70^Zn associated with the cells were determined by ICP-MS. As presented in **Figure 4B**, our calculation showed that 6.4±3.9% of the endogenous ^64^Zn was expelled from the cells during the 30 min transport assay. The calculations are detailed in *Experimental procedures*. Given that the imported zinc ions rapidly exchange with a single pool of the endogenous zinc (**Figure 4A**), it can be deduced that the same percentage of ^70^Zn imported during the assay has been expelled from the cells through the same pathway as the endogenous ^64^Zn. This result indicates that a small fraction of zinc ions from the endogenous zinc pool (^64^Zn) and those from the extracellular space (mainly ^70^Zn) are exported from the cells during the transport assay, and therefore the hZIP4 activity (determined by the imported ^70^Zn) is indeed underestimated. Given that only ∼6% of the influxed zinc was exported during the assay, we conclude that zinc efflux would not significantly affect the transport kinetic study of hZIP4 under the current experimental conditions. Consistently, a fit of the data in **Figure 4A** indicates that an approximate half life for zinc loss from the cells was 3 hours (i.e. 164 min), providing further support for this conclusion. It has been reported that mammalian cells handle light and heavy zinc isotopes differently, resulting in a 0.2-0.4‰ difference in their transport (80), which is two orders of magnitude smaller than the ∼6% zinc efflux determined in this work. Therefore, the kinetic isotope effect may have little effect on data interpretation. Nevertheless, our data also suggest that care should be taken in the experimental design to minimize the effect of substrate efflux on the activity measurement of an importer.

**Figure 4.**
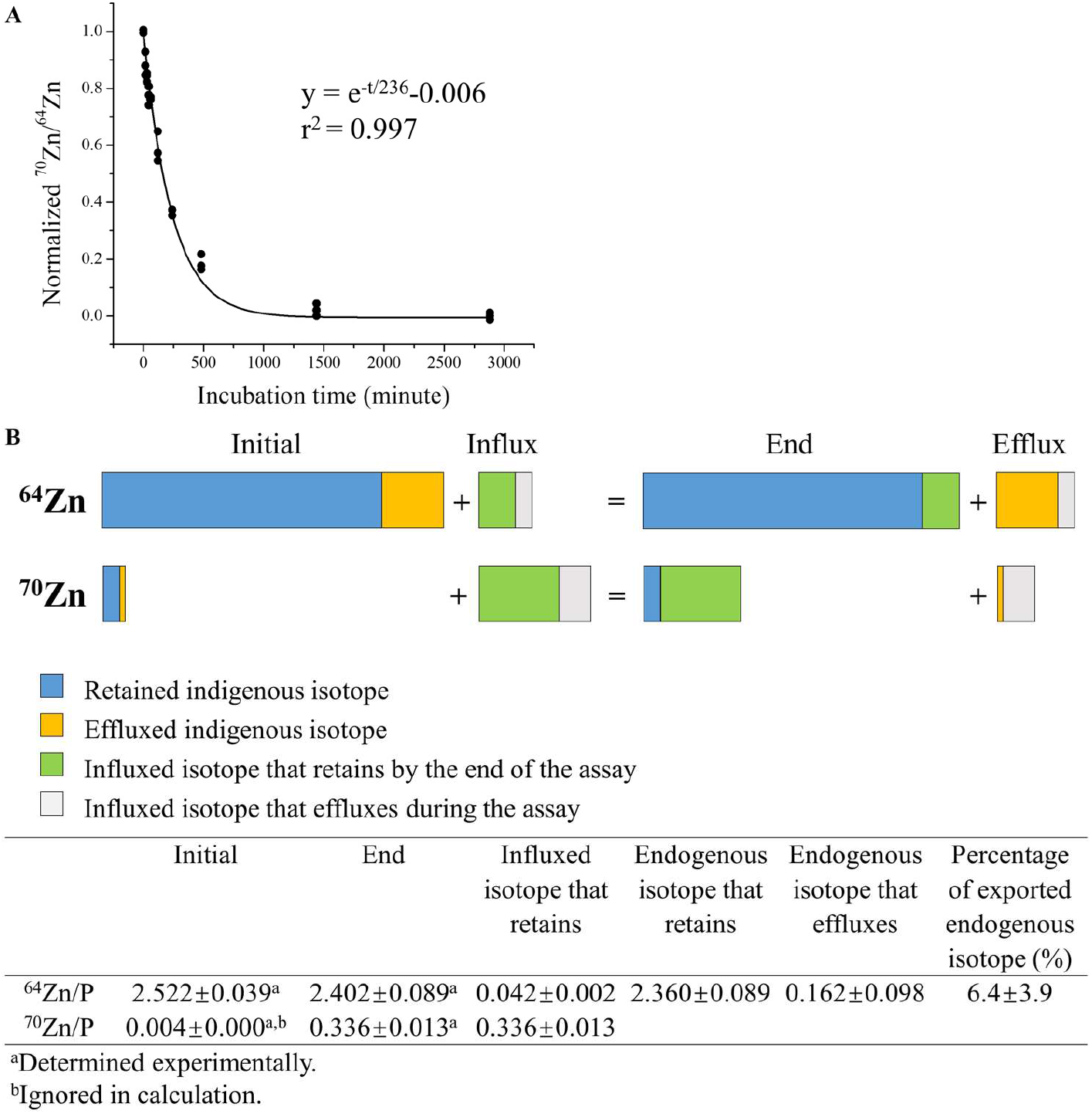
Evaluation of zinc efflux during the transport assay. (**A**) Time course of ^70^Zn replacement by ^64^Zn. Cells growing in the regular culture media with naturally present zinc isotopes were incubated with the ^64^Zn-enriched culture media. At the indicated time points, cells were digested for ICP-MS analysis. The ^70^Zn/^64^Zn count ratios were normalized and plotted against the reaction time, and the curve was fitted using the one-phase decay model. (**B**) Estimation of zinc efflux during the transport assay. The hZIP4-expressing cells growing in the ^64^Zn-enriched culture media were washed and then incubated with the ^70^Zn-enriched culture media for 30 minutes. The ^64^Zn/P and ^70^Zn/P count ratios were determined by ICP-MS before and after the transport assay. *Upper*: The diagram illustrating the changes of ^64^Zn and ^70^Zn due to influx and efflux during the assay. *Lower*: Calculation of zinc efflux in one of three independent experiments in which similar results were obtained. Six replicates were performed for each condition. The calculations are described in detail in *Experimental procedures*.

Collectively, it is very unlikely that the determined slow transport rate (0.08-0.2 s^-1^) is due to a significant zinc efflux during the transport assay. We speculate that in order to acquire an adequate amount of zinc using a slow zinc transporter like ZIP4, cells must express a large number of transporter molecules at the cell surface, thus allowing the mechanisms that regulate the cell surface expression level of the transporter to efficiently fine tune the overall zinc transport capacity at the plasma membrane. Whether endogenously expressed hZIP4, other than hZIP4 overexpressed in HEK293T cells, exhibits a similar transport rate and whether other mammalian ZIPs are also slow transporters warrant future investigation.

### LA-ICP-MS assisted zinc transport assay with an increased throughput

It takes approximately three minutes to analyze one sample by ICP-MS in the liquid mode. A higher throughput is required for many applications, including inhibitor screening for drug discovery and directed evolution for transporter engineering. We explored the possibility of increasing the speed of isotope measurement by using LA-ICP-MS. For this purpose, cells were lysed after the ^70^Zn-based transport assay, and the cell lysate was dropped onto a siliconized glass slide and dried in oven (**Figure 5A)**. On a standard glass slide (26 mm x 75 mm), up to 36 drops can be manually applied at a volume of 5 μl per drop. The ^70^Zn/P count ratio of each drop was then measured by LA-ICP-MS equipped with a time-of-flight (TOF) detector. Because the ^70^Zn/P ratio remains nearly constant at different locations within one drop (**Figure S6**), each drop was scanned by five parallel lines across the drop, rather than the entire area of the drop, to speed up the process. Accordingly, it took less than half a minute to scan one drop, reducing the analysis time by a factor of six when compared with the assays conducted in the liquid mode. Although the absolute amount of ^70^Zn associated with the cells are not established under the current protocol, a dose curve of hZIP4-mediated zinc transport can be generated by using the determined ^70^Zn/P ratios. As shown in **Figure 5B**, the ^70^Zn/P ratios in the hZIP4-expressing cells are significantly higher than those in the cells transfected with the empty vector, and subtraction of the latter from the former yielded a dose curve similar to that generated by using the data collected by ICP-MS in the liquid mode (**Figure 1**). Curve fitting using the Hill model resulted in an apparent *K*_M_ of 2.3±0.3 μM (n=3) with the Hill coefficient of ∼1 (**Figure 5A**), consistent with values obtained in prior studies (54,56) and in the transport assay described above (**Figure 1**). These results demonstrate that this LA-ICP-TOF-MS assisted approach significantly increases the throughput without compromising data quality. We expect the throughput of this approach to further increase significantly with optimization of automation of cell handling, slide preparation, and slide scanning strategy.

**Figure 5.**
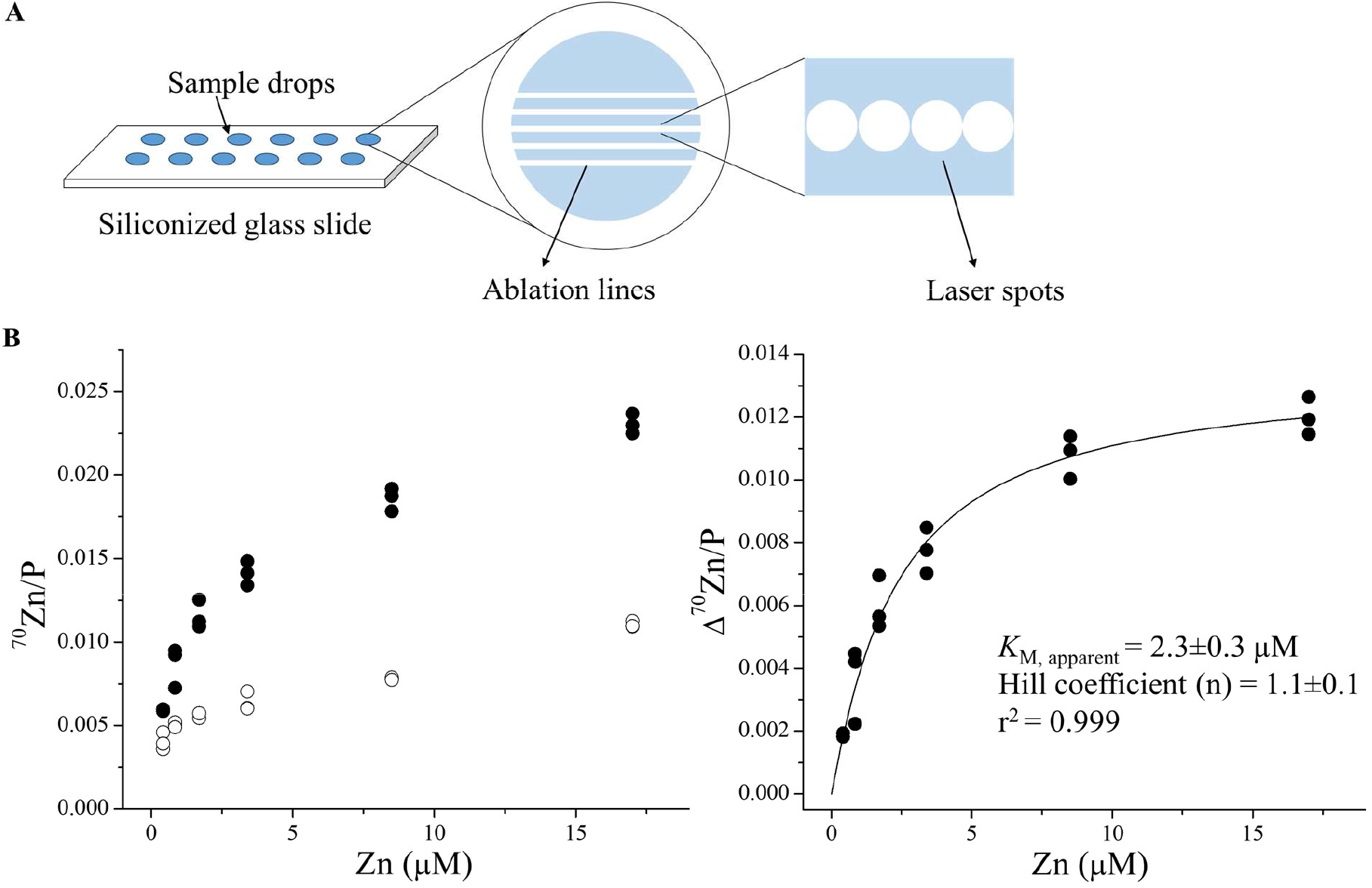
Kinetic study of hZIP4 with increased throughput by using LA-ICP-MS. (**A**) Cartoon illustration of a siliconized glass slide loaded with cell lysate samples for LA-ICP-MS detection. To increase the detection speed, five parallel lines (composed of continuous contacting spots) were ablated across each sample drop. (**B**) Dose curve of hZIP4-mediated ^70^Zn transport using LA-ICP-TOF-MS to detect ^70^Zn and ^31^P. *Left*: The ^70^Zn/P count ratios of the cells with (solid circle) and without (open circle) expressing hZIP4 at the indicated concentrations of total zinc in the extracellular space. *Right*: Kinetic study of hZIP4. The activity of hZIP4 was calculated by subtracting the ^70^Zn/P count ratio of the cells transfected with an empty vector from that of the cells expressing hZIP4. The curve was fitted using the Hill model. Each point represents the data of the cells in one subwell. For each condition, three replicates were performed. The extracellular zinc consists of ^70^Zn (with an enrichment of 72%) and other zinc isotopes. The shown data are from one of three independent experiments in which similar results were obtained.

## Conclusion

In this work, we applied a modified ^70^Zn-based approach to study the transport kinetics of hZIP4, a representative ZIP metal transporter and a promising oncology drug target. Our data showed that the low-abundance stable isotope ^70^Zn can adequately replace the radioactive ^65^Zn in the cell-based transport assay. This approach, combined with quantification of the surface expression level of hZIP4, allowed us to estimate the apparent turnover number of hZIP4 to be in the range of 0.08-0.2 s^-1^, indicating that hZIP4 is best described at this time as a carrier, rather than an ion channel. Re-examination of the role of the hZIP4-ECD reinforced the notion that the HRD subdomain is responsible for most of the function of the ECD in facilitating zinc transport. Estimation of zinc efflux during the transport assay showed that zinc efflux did not significantly affect the analysis of zinc influx. We also demonstrated that the throughput of the ^70^Zn-based approach can be significantly increased when LA-ICP-TOF-MS was used to study the samples loaded on glass slides. This approach may be developed into an assay suitable for drug discovery targeting other ZIPs and zinc transporters from other metal transporter families. Transport assay of the biological metals with more than one stable isotopes, such as iron (^54^Fe, ^56^Fe, ^57^Fe, and ^58^Fe), nickel (^58^Ni, ^60^Ni, ^61^Ni, ^62^Ni, and ^64^Ni), and copper (^63^Cu and ^65^Cu), can also be performed using the ICP-MS-based approaches presented in this work.

## Acknowledgments

We thank Tianqi Wang in the Hu laboratory for providing the purified FLAG-hZIP4-HA. We thank Dr. Aaron Sue in O’Halloran laboratory for providing assistance in the ICP-MS and LA-ICP-MS experiments. We thank Dr. Benjamin Orlando in the Department of Biochemistry and Molecular Biology at Michigan State University for helping us generate the structural models of mouse ZIP4 using AlphaFold 2.0.

## Author contributions

J. H., Y.J., K.M., and T.V.O. conceived the project and designed the experiments, Y.J. and K.M. conducted the experiments; Y.J., K.M., T.V.O., and J. H. analyzed the data and wrote the manuscript.

## Funding and additional information

This work is supported by National Institutes of Health GM129004 and GM140931 (to J. H.) and GM135018, GM115848, and GM038784 (to T.V.O.).

The content is solely the responsibility of the authors and does not necessarily represent the official views of the National Institutes of Health.

## Experimental procedures

### Gene, plasmids, and reagents

The complementary DNA of human ZIP4 (GenBank access number: BC062625) from Mammalian Gene Collection were purchased from GE Healthcare. The hZIP4 coding sequence was inserted into a modified pEGFP-N1 vector (Clontech) in which the downstream EGFP gene was deleted and an HA tag was added at the C-terminus. ZIP4 variants (ΔHRD-hZIP4-HA and ΔECD-hZIP4-HA) were generated as previously described (54). All mutations were verified by DNA sequencing. For the hZIP4 construct for purification, a FLAG tag is added immediately after the signal peptide to generate the final FLAG-hZIP4-HA construct (75). ^64^ZnO and ^70^ZnO were purchased from American Elements (Product# ZN-OX-01-ISO.064I, Lot#1871511028-400 and ZN-OX-01-ISO.070I, Lot#1871511028-401, respectively). 30 mg of zinc oxide powder was dissolved in 5 ml of 1 M HCl and then diluted with ddH2O to make the stock solution at the concentration of 50 mM. The ^64^Zn sample was certified as 98% abundance while ^70^Zn isotope in the other sample corresponded to an abundance of 72%. Other reagents were purchased from Sigma-Aldrich or Fisher Scientific.

### Cell culture, transfection, and Western blot

Human embryonic kidney cells (HEK293T, ATCC, #CRL-3216) were cultured in Dulbecco’s modified eagle medium (DMEM, Thermo Fisher Scientific, Invitrogen, Cat#11965092) supplemented with 10% (v/v) fetal bovine serum (FBS, Thermo Fisher Scientific, Invitrogen, Cat#10082147) and Antibiotic-Antimycotic solution (Thermo Fisher Scientific, Invitrogen, Cat# 15240062) at 5% CO_2_ and 37°C. The polystyrene 24-well trays (Alkali Scientific, Cat#TPN1024) were treated with 300 μl Poly-D-Lysine (Corning, Cat#354210) for each well overnight, after which cells were seeded in the DMEM plus 10% FBS medium. After 16 hours of incubation cells were transfected with 0.8 μg DNA/well using lipofectamine 2000 (Thermo Fisher Scientific, Invitrogen, Cat# 11668019) in DMEM plus 10% FBS.

For Western blot, samples were mixed with the SDS sample loading buffer and heated at 96 °C for 10 min before loading on SDS-PAGE gel. The proteins separated by SDS-PAGE were transferred to PVDF membranes (Millipore, Cat#PVH00010). After blocking with 5% (w/v) nonfat dry milk, the membranes were incubated with mouse anti-HA antibody (Thermo Fisher Scientific, Cat# 26183) or rabbit anti-β-actin (Cell Signaling, Cat# 4970 S) at 4 °C overnight, which were detected with HRP-conjugated horse anti-mouse immunoglobulin-G antibody at 1:5000 dilution (Cell Signaling Technology, Cat# 7076) or goat anti-rabbit immunoglobulin-G antibody at 1:3000 dilution (Cell Signaling Technology, Cat# 7074) respectively using the chemiluminescence reagent (VWR, Cat#RPN2232). The images of the blots were taken using a Bio-Rad ChemiDoc Imaging System.

### Treatment of the culture media with Chelex-100

100 ml of the culture media (DMEM+10% FBS) was mixed with 5 g Chelex-100 resin (Sigma-Aldrich, Cat# C7901) in a column and placed on a gyratory rocker overnight. The resin can be regenerated for further treatment after following process: washed by one column volume of water, incubate with one column volume of 1 M hydrochloric acid for one hour, wash with one column volume of water, incubate with one column volume of 1 M sodium hydroxide for one hour, wash with two column volumes of water.

### Zinc transport assay

Twenty hours post transfection, cells were washed with the washing buffer (10 mM HEPES, 142 mM NaCl, 5 mM KCl, 10 mM glucose, pH 7.3) followed by incubation with Chelex-treated culture media. ^70^Zn was added to initiate the transport. In the experiments shown in Figures 1A, 1B and 4B, 5 μM of ^70^Zn was added. In the experiments shown in Figures 2 and 3, 12 μM of ^70^Zn (about 5-10 times of the apparent *K*_M_) was added. After incubation at 37°C for 30 min, the plates were put on ice and an equal volume of the ice-cold washing buffer containing 1 mM EDTA was added to the cells to terminate metal uptake, followed by three times of washing with ice-cold washing buffer.

### Estimation of cell surface expression level by biotinylation

Cell surface protein biotinylation and isolation was done with the cell surface protein biotinylation and lsolation kit (Thermoscientific, Cat#A44390). 20 hours post transfection, the cells from six wells in a 24-well tray were washed by phosphate-buffered saline (PBS, 200 μl per well) and subjected to amine labelling of the membrane proteins at cell surface with Sulfo-NHS-SS-Biotin in PBS (200 μl per well) as described in the manual. After incubation at room temperature for 10 minutes, the labelling solution was removed and cells were washed with the ice-cold tris(hydroxymethyl)aminomethane-buffered saline (TBS, 200 μl per well) to terminate the reaction. Cells were then harvested in TBS (200 μl per well) and lysed. The lysate was applied to the NeutrAvidin agarose column to purify the labeled proteins. The labelled proteins were collected in the elution buffer containing 10 mM DTT. The samples were then mixed with 2xSDS loading buffer with a 1:1 volume ratio. hZIP4 with an N-FLAG tag and C-HA tag (FLAG-hZIP4-HA) was purified as reported (75) and used as the standard to generate a standard curve. The samples from cells and the standards were subjected to Western blot.

### Estimation of cell surface expression level by surface bound antibody

Twenty hours post transfection, cell culture media was removed and cells were washed with Dulbecco’s phosphate-buffered saline (DPBS). Cells were then fixed with 4% formaldehyde for 10 minutes at room temperature, followed by three times of washing with DPBS to remove formaldehyde. To detect the HA-tagged hZIP4 expressed at cell surface, cells were incubated in DPBS solution with 5% BSA (Sigma-Aldrich, Cat# A2153) and 3 μg/ml anti-HA antibody (Invitrogen, Cat# 26183) for one hour and a half at room temperature. The solution was then removed and the cells were washed five times with DPBS. Each time the plate was put on a gyratory rocker for 5 minutes of gentle shaking. The sample was subjected to Western blot together with the known amount of anti-HA antibody. The latter was used to generate a standard curve to estimate the cell surface bound anti-HA antibody. To detect the cell surface bound anti-HA antibodies, the HRP-conjugated horse anti-mouse immunoglobulin-G antibody at 1:5000 dilution (Cell Signaling Technology, Cat# 7076) was directly applied to the PVDF membrane in the Western blot experiment, followed by signal visualization using the chemiluminescence reagent (VWR, Cat#RPN2232).

### Zinc isotope replacement experiment and estimation of zinc efflux

HEK293T cells were seeded into poly-D-lysine treated polystyrene 24-well plates as described previously and allowed to grow in the regular culture media (DMEM plus 10% FBS) for 16 hours. The culture media were then removed as much as possible and the metal supplemented ^64^Zn-enriched culture media was added. The metal supplemented ^64^Zn-enriched culture media is the Chelex-treated culture media (DMEM plus 10% FBS) supplemented with Mg, Ca, Cu, and ^64^Zn to restore these metals back to the levels in the untreated culture media.. The calculation was based on the ICP-MS data in **Table 1** At the designed time points, the ^64^Zn enriched culture media was removed and cells were washed with the washing buffer for three times. The cells were digested in nitric acid for ICP-MS analysis.

### Determination of zinc efflux during the transport assay

After being cultured in the ^64^Zn enriched culture media for three passages in a poly-D-lysine treated polystyrene 24-well plate, HEK293T cells were transfected with the plasmid encoding hZIP4. 20 hours post transfection, the cells were treated with or without the ^70^Zn-enriched culture media containing 5 μM ^70^Zn. Six replicates from different wells were included in each group (the treated cells or the untreated cells). After incubation at 37°C for 30 minutes, the cells in both groups were treated in the same way as described in the section of zinc transport assay. ^64^Zn and ^70^Zn associated with the cells were quantified by ICP-MS and calibrated with ^31^P in the same sample by using the Zn/P count ratio. A cartoon illustration explaining the following equations is shown in **Figure 4B**. The efflux of ^64^Zn during the transport assay was determined accordingly.

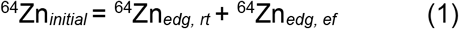

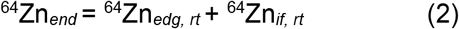

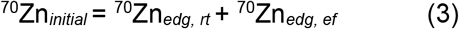

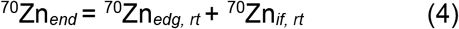

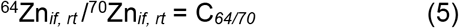

where the subscripts *initial* and *end* represent states of cells before and after the assay, respectively. *edg* represents the endogenous zinc; *rt* represents the zinc retained after the assay; *ef* represents the zinc exported during the assay; *if* represents the zinc imported during the assay.

^64^Zn_*initial*_ and ^70^Zn_*initial*_ were determined in the untreated cells, and ^64^Zn_*end*_ and ^70^Zn_*end*_ were determined in the treated cells. C_*64/70*_ refers to the ^64^Zn/^70^Zn molar ratio in the ^70^Zn stock solution, which was experimentally determined to be 0.123. As shown in the dataset in **Figure 4B**, ^70^Zn_*initial*_ is two orders of magnitude smaller than ^70^Zn_*end*_, which is because the cells were grown in the ^64^Zn-enriched culture media. Since ^70^Zn_*edg, ef*_ is a portion of ^70^Zn_*initial*_, it can be ignored in (4), yielding

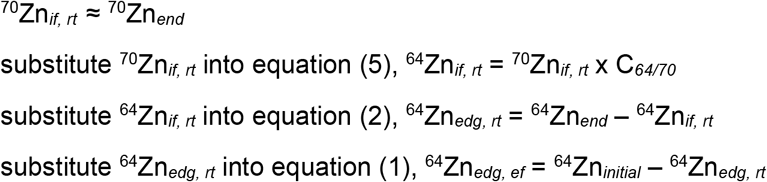

Accordingly, percentage of endogenous ^64^Zn efflux = ^64^Zn_*edg, ef*_ / ^64^Zn_*initial*_ x 100%.

### ICP-MS

All standards, blanks, and cell samples were prepared using trace metal grade nitric acid (70%, Fisher chemical, Cat# A509P212), ultrapure water (18.2 MΩ·cm @ 25 °C), and metal free polypropylene conical tubes (15 and 50 mL, Labcon, Petaluma, CA, USA). For cell samples in polystyrene 24-well cell culture plates, 200 μl of 70% trace nitric acid was added to allow for initial sample digestion. Following digestion, 150 μl of the digested product was transferred into metal free 15 mL conical tubes. For liquid samples, 50 μl of liquid samples were added to metal free conical tubes followed by addition of 150 μl of 70% trace nitric acid. All cell and liquid samples were then incubated at 65°C in a water bath for one hour followed by dilution to 5 ml using ultrapure water. These completed ICP-MS samples were then analyzed using the Agilent 8900 Triple Quadrupole ICP-MS (Agilent, Santa Clara, CA, USA) equipped with the Agilent SPS 4 Autosampler, integrate sample introduction system (ISiS), x-lens, and micromist nebulizer. Daily tuning of the instrument was accomplished using manufacturer supplied tuning solution containing Li, Co, Y, Ce, and Tl. Global tune optimization was based on optimizing intensities for ^7^Li, ^89^Y, and ^205^Tl while minimizing oxides (^140^Ce^16^O/^140^Ce < 1.5%) and doubly charged species (^140^Ce++/^140^Ce+ < 2%). Following global instrument tuning, gas mode tuning was accomplished using the same manufacturer supplied tuning solution in KED mode (using 100% UHP He, Airgas). Specifically, intensities for ^59^Co, ^89^Y, and ^205^Tl were maximized while minimizing oxides (^140^Ce^16^O/^140^Ce < 0.5%) and doubly charged species (^140^Ce++/^140^Ce+ < 1.5%) with short term RSDs < 3.5%. ICP-MS standards were prepared from a stock solution of NWU-16 multi-element standard (Inorganic Ventures, Christiansburg, VA, USA) that contains As, Ca, Cd, Co, Cr, Cu, Fe, K, Mg, Mn, Ni, Se, V, Zn that were diluted with 3% (v/v) trace nitric acid in ultrapure water to a final element concentration of 1000, 500, 250, 125, 62.5, 31.25, and 0 (blank) ng/g standard. Internal standardization was accomplished inline using the ISIS valve and a 200 ng/g internal standard solution in 3% (v/v) trace nitric acid in ultrapure water consisting of Bi, In, ^6^Li, Sc, Tb, and Y (IV-ICPMS-71D, Inorganic Ventures, Christiansburg, VA, USA). The isotopes selected for analysis were ^31^P, ^64^Zn, ^66^Zn, ^67^Zn, ^68^Zn, ^70^Zn and ^6^Li, ^45^Sc, and ^89^Y for internal standardization. Continuing calibration blanks (CCBs) were run every 10 samples and a continuing calibration verification standard was analyzed at the end of every run for a 90-110% recovery.

### LA-ICP-TOF-MS

Cell samples were lysed with 0.25% Triton X-100 plus 10% Coomassie blue. 5 μl of cell lysate was carefully dropped onto a siliconized glass slide which was then transferred into a 200°C oven to let the droplets dry completely. To siliconize glass slides, we used 25 x 75 x 1.0 mm microscope slides (Alkali Scientific, Cat# SM2551) that were washed thoroughly with hand soap and water. After leaving on bench top for drying, the slides were placed in a beaker to which siliconization solution (Supelco, Cat# 85126) was added, and the slides were submerged covered for 30 minutes at room temperature (about 25°C) in a hood. The glass slides were then carefully removed from the beaker and dried overnight at room temperature. The slides were then rinsed briefly with ultrapure deionized water and placed in an oven (45°C) overnight. The slides need to be preheated before samples are added to minimize the coffee stain effect and for sufficient drying of the droplet.

Glass slides were loaded into a Bioimage 266 nm laser ablation system (Elemental Scientific Lasers, Bozeman, MT, USA) which is equipped with an ultra-fast low dispersion TwoVol3 ablation chamber and a dual concentric injector (DCI3) and is coupled to an icpTOF S2 (TOFWERK AG, Thun, Switzerland) ICP-TOF-MS. Daily tuning of the LA-ICP-TOF-MS settings was performed using NIST SRM612 glass certified reference material (National Institute for Standards and Technology, Gaithersburg, MD, USA). Optimization for torch alignment, lens voltages, and nebulizer gas flow was based on high intensities for ^140^Ce and ^55^Mn while maintaining low oxide formation based on the ^232^Th^16^O+/^232^Th+ ratio (< 0.5).

Sample loaded glass slides were tested on the LA system using multiple laser powers and spot sizes to determine optimum parameters for ablation. Due to the sample dye, we needed higher LA power to ablate through the sample with enough energy density while still minimizing ablation of glass. The optimized laser parameters were 80% laser power (9.5-10.5 J/cm^2^ laser fluence, 0.1-0.15 mJ sample energy) with a 40 μm spot size (circular) and repetition rate of 50 Hz (single pulse response was tested daily with values between 4-5 ms) with no overlap between the adjacent laser spots. Ablation of the glass slide was analyzed using ^27^Al and ^140^Ce elemental maps to ensure minimal glass ablation but complete sample ablation. Following ablation of the entire droplet and concurrent analysis (**Figure S6**), we sped up the process by ablating five parallel lines across each droplet with 120 μm spacing between two adjacent lines. This provided sufficient data points for statistical analysis with minimal differences between analyses (**Figure S6**). Because of the coffee stain effect, cell lysate was not evenly distributed within a sample droplet. For each ablation line, the ^31^P counts were used to identify and exclude the data from the ablation points where the cell lysate content is very low. Then, the counts of ^31^P and ^70^Zn from the five ablation lines for each sample droplet were summed up respectively and used to calculate the ^70^Zn/^31^P count ratio. The instrument parameters for all MS experiments are listed in **Table S1**.

### Statistics

We assumed a normal distribution of the samples and significant difference were examined using Student’s *t* test (two tailed). Uncertainties are reported as standard deviation or standard error of the mean, as indicated.

