## Supplementary Information for "Determination of metal ion transport rate of human ZIP4 using stable zinc isotopes"

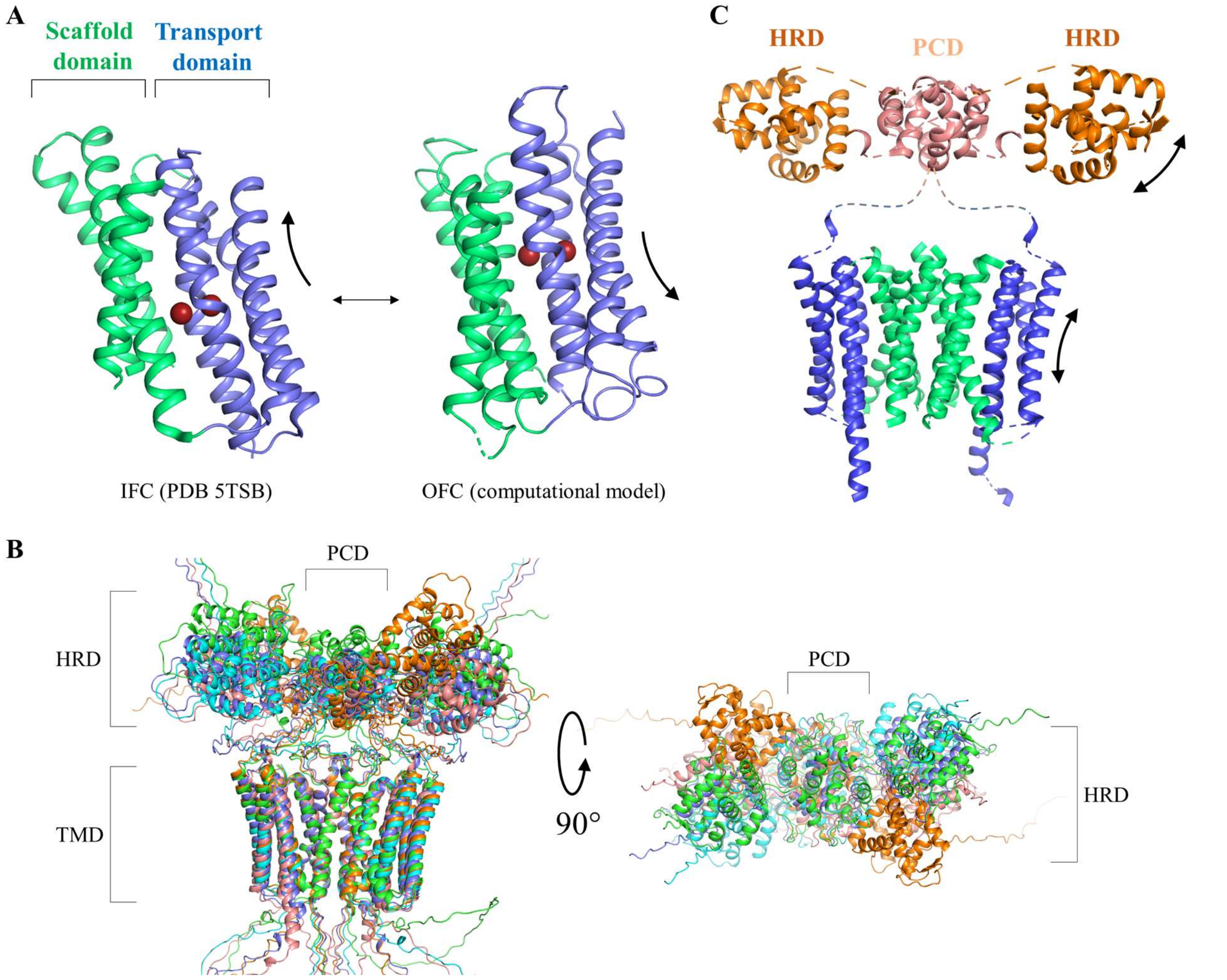


**Scheme S1.** Proposed elevator-type transport mechanism. (**A**) The elevator motion of BbZIP. The interconversion of the inward-facing and outward-facing conformations is achieved by vertical sliding of the transport domain (blue) against the static scaffold domain (green) that is involved in dimerization (not shown). The computational model for the outward-facing conformation was reported in ref 40. (**B**) Five AlphaFold 2.0 predicted structures of mouse ZIP4 dimer in the side (*left*) and top (*right*) views. The transmembrane domains (TMD) are structurally superimposed to reveal the flexibility of the HRD and PCD domains. (**C**) Putative interactions between the HRD (orange) and the transport domain (blue). The TMD is colored as for BbZIP in (A). For clarity, loops have been trimmed or shown as dashed lines. Considering that the elevator motion of the transport domain and the flexibility of the HRD domain, as indicated by the curved arrows, it is plausible that the interaction of the HRD with the transport domain may promote the formation and stabilization of the outward-facing conformation and thereby facilitate zinc transport.

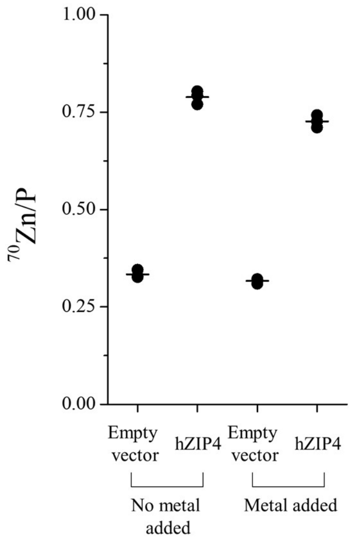


**Figure S1.** Comparison of hZIP4 activity in the Chelex-treated culture media with and without supplementing Mg, Ca, and Cu. These metal elements were found to be removed by Chelex-100 (**Table 1**).

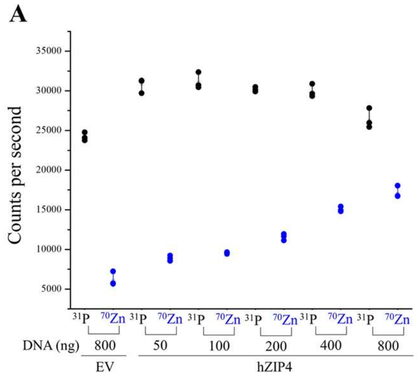


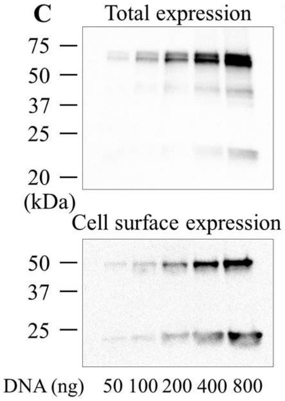


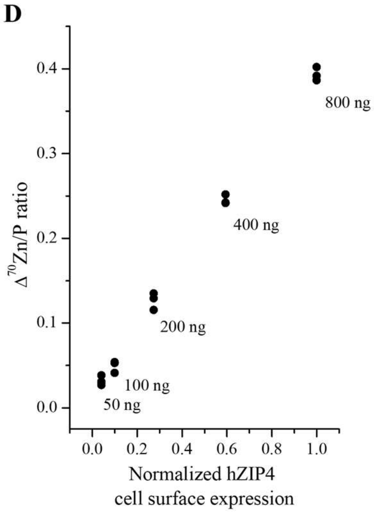


**B**

| Plasmid DNA (ng) | EV | 50 | 100 | 200 | 400 | 800 |
|---|---|---|---|---|---|---|
| $^{70}$Zn/P | 0.257 | 0.289 | 0.307 | 0.384 | 0.502 | 0.651 |
| $^{70}$Zn/P after background subtraction | | 0.032 | 0.049 | 0.126 | 0.245 | 0.393 |

**Figure S2.** Correlation between the zinc transport activity of hZIP4-HA and its cell surface expression level in HEK293T cells. (**A**) Counts (per second, detected in ICP-MS) of $^{31}$P and $^{70}$Zn in the HEK293T cells transfected with empty vector (EV, 800 ng) or the indicated amounts of plasmid DNA encoding hZIP4-HA (50-800 ng). The error bars represent the standard deviation of the mean. (**B**) $^{70}$Zn/P count ratios before and after subtracting the background. (**C**) Total and cell surface expression of hZIP4. Cells in each well of a 24-well plate were transfected with the indicated amounts of the plasmid DNA with a fixed ratio of 400 ng DNA per µL of Lipofectamine 2000. The total expression of hZIP4 was detected in Western blot using an anti-HA antibody, and the cell surface expression of hZIP4 was determined by the surface bound anti-HA antibody, which was detected in Western blot using an HRP-conjugated horse anti-mouse IgG antibody. (**D**) Correlation of the $^{70}$Zn/P ratios (after background subtraction) and the cell surface expression levels of hZIP4.

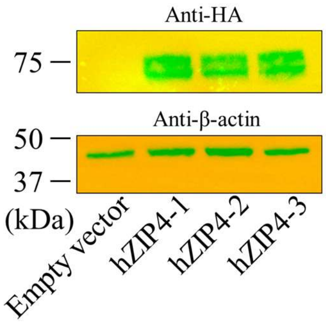


**Figure S3.** Comparison of hZIP4 transient expression in the cells grown in different wells in the same experiment. Western blot was performed to detect the HA-tagged hZIP4 and β-actin.

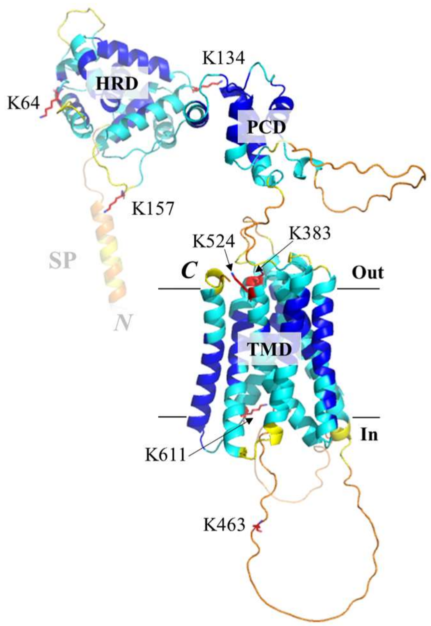


**Figure S4.** Lysine residues in hZIP4. The AlphaFold predicted structure of hZIP4 (UniProt ID: Q6P5W5) is shown in cartoon mode and colored with the confidence score (pLDDT). hZIP4 consists of the ECD and the TMD, and the former is composed of the very N-terminal HRD subdomain and the PCD subdomain. All the seven lysine residues are shown in stick mode, labeled, and colored in red, out of which five are exposed to the extracellular side – three are fully exposed in the HRD and two appear to be partially buried in the TMD. SP: signal peptide.

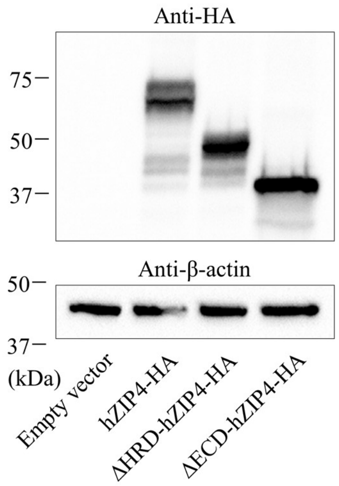


**Figure S5.** Comparison of total expression of hZIP4 and the truncated variants. Western blot was performed to detect the HA-tagged proteins and β-actin.

A

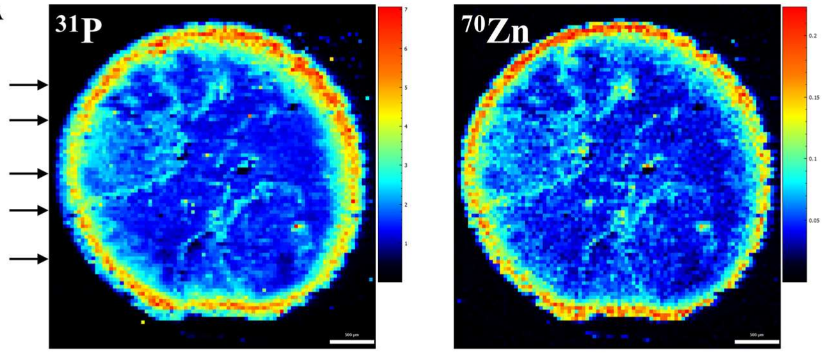


| | Whole drop | Sum of the five lines |
|---|---|---|
| $^{31}$P | 154.1289 | 9.3553 |
| $^{70}$Zn | 4.7142 | 0.2721 |
| $^{70}$Zn/$^{31}$P | 0.0306 | 0.0291 |

B

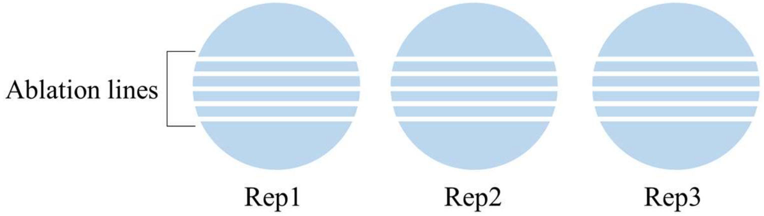


C

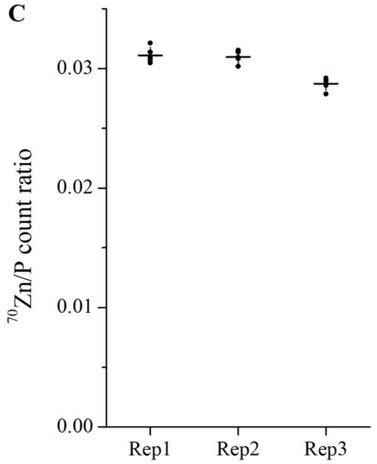


**Figure S6.** Comparison of the $^{70}$Zn/P count ratios at different locations within sample drops. **(A)** Imaging of $^{31}$P and $^{70}$Zn of a drop of cell lysate on glass slide by LA-ICP-MS. The arrows indicate the rough positions of the five ablation lines randomly selected for analysis. The scale bars indicate 500 µm. The table shows that the determined $^{70}$Zn/$^{31}$P count ratios for the whole drop and the sum of the five lines are nearly identical. (**B**) Cartoon illustration of ablation of three replicates. The white lines represent the ablation lines. For each drop, ablation was conducted along five parallel lines with a distance of 120 micrometers between the adjacent lines. (**C**) Comparison of the $^{70}$Zn/P count ratios of the five lines within the and between the sample drops. The shown data are the three replicates of the sample for which the hZIP4-expressing HEK293 cells were treated with the $^{70}$Zn-enriched culture media containing 17 µM total zinc. Each dot represents the $^{70}$Zn/P count ratio of one ablation line, which was determined by LA-ICP-MS. The relative SD for the five lines in each replicate is approximately 2%, indicating that the $^{70}$Zn/P count ratios at different locations in one sample drop are highly consistent.

**Table S1.** Parameters for the experiments of ICP-MS and LA-ICP-MS.

| ***ICP-MS (Tofwerk S2)*** | | |
|---|---|---|
| **Parameter** | Unit | Value |
| **RF Power** | W | 1550 |
| **Sampling Depth** | mm | 4.9 |
| **Cone Material** | Nickel | - |
| **Cone Insert (STD)** | mm | 3.5 |
| **Plasma Gas Flow** | L/min | 14.0 |
| **Auxillary Gas Flow** | L/min | 8.0 |
| **Nebulizer Gas Flow** | L/min | 0.95-1.01 |
| **Measurement Mode** | CCT Mode | - |
| **CCT Gas Flow (100% He)** | mL/min | 5 |
| **CCT Focus lens** | V | -19.5 |
| **CCT Entry Lens** | V | -180 |
| **CCT Mass** | V | 250 |
| **CCT Bias** | V | 10 |
| **CCT Exit Lens** | V | -200 |
| ***Time-of-Flight (Tofwerk S2)*** | | |
| **Parameter** | Unit | Value |
| **m/z range** | amu | 14 - 256 |
| **Resolution** | m/Δm | 1000 |
| **ODG Settings** | ms | 24 |
| **Notch Bias** | V | -80 |
| **Notch (40 amu)** | V | 3 |
| **Notch (36 amu)** | V | 1 |
| **Notch (28 amu)** | V | 2 |
| **Notch (15.8 amu)** | V | 3 |
| ***Laser Ablation (ESL Bioimage 266 nm)*** | | |
| **Parameter** | Unit | Value |
| **Spot Size** | µm | 40 |
| **Interline Distance (y)** | µm | 120 |
| **Overlap (x)** | µm | 0 |
| **Repetition Rate** | Hz | 50 |
| **Laser Power** | % | 80 |
| **Laser Fluence** | $J/cm^2$ | 9.5-10.5 |
| **Sample Energy** | mJ | 0.1-0.15 |
| **Imaging Cup Flow Rate (He)** | mL/min | 200 |
| **Imaging Chamber Flow Rate (He)** | mL/min | 250 |
| **PEEK Tubing I.D.** | mm | 0.75 |
